# Role of *ugt* genes in detoxification and glycosylation of 1-hydroxy phenazine (1-HP) in *Caenorhabditis elegans*

**DOI:** 10.1101/2023.11.21.568030

**Authors:** Muhammad Zaka Asif, Kelsey A. Nocilla, Li T. Ngo, Man K. Shah, Yosef Smadi, Zaki A. Hafeez, Michael Parnes, Allie Manson, John Glushka, Franklin E. Leach, Arthur S. Edison

## Abstract

*Caenorhabditis elegans* is an ideal model organism to study the xenobiotic detoxification pathways of various natural and synthetic toxins. One toxin shown to cause death in *C. elegans* is 1-hydroxyphenazine (1-HP), a molecule produced by the bacterium *Pseudomonas aeruginosa.* We previously showed that the median lethal dose (LD50) for 1-HP in *C elegans* is 179 μM in PD1074 and between 150-200 μM in N2 (*C. elegans* lab strain). We also showed that *C. elegans* detoxifies 1-HP by glycosylation by adding one, two, or three glucose molecules in N2 worms. This study tested whether UDP-glycosyltransferase (*ugt)* genes play a role in 1-HP detoxification. We show that *ugt-23* and *ugt-*49 knockout mutants are more sensitive to 1-HP. Our data also show that *ugt-23* knockout mutants produce reduced amounts of the trisaccharide sugars, while the *ugt-49* knockout mutants produce reduced amounts of all 1-HP derivatives except for the glucopyranosyl product. We have also characterized the structure of the trisaccharide sugar phenazine structures made by *C. elegans* and show that one of the sugar modifications contains an N-acetylglucosamine (GlcNAc) in place of glucose. This implies broad specificity regarding UGT function and the role of genes other than *ogt-1* in adding GlcNAc, at least in small-molecule detoxification.

---

*C. elegans* are bacterivores found in soil and decaying organic matter.^[1, 2]^ As they feed on bacteria in their environment, they are exposed to numerous pathogens and xenobiotics.^[3]^ To combat exposure, worms have developed three main strategies for defense. One is avoidance, where they can sense potentially hostile environments and avoid going to them.^[4, 5]^ Another is the presence of a strong cuticle and pharyngeal grinder to physically prevent pathogens from entering the worm.^[6]^ Finally, if pathogens can enter the worm, several mechanisms are activated, constituting the innate immune response.

Along with pathogen response, the *C. elegans* innate immune system is activated upon xenobiotic exposure. Xenobiotics are defined as substances foreign to a body or ecological system. In nature, *C. elegans* feed on various bacteria, many of which produce toxic compounds to the worms. As a result, *C. elegans* have developed a wide array of detoxification enzymes.^[7]^ Xenobiotic metabolism is canonically divided into three phases.^[8]^ Phase I is the addition of reactive moieties, such as hydroxyl groups, to the parent xenobiotic. Phase II is the conjugation of either the phase I modified or parent xenobiotic to a large, water-soluble molecule to facilitate excretion. Phase III is the transport of these metabolized compounds out of the cell.^[9]^ In this study, we focus on steps involved with the phase II detoxification of one such xenobiotic: 1-hydroxyphenazine (1-HP).

1-Hydroxyphenazine (1-HP) is a small molecule produced by many *Pseudomonas* species, including *Pseudomonas aeruginosa.*^[10–14]^ 1-HP is one of three related metabolites, along with pyocyanin and phenazine-1-carboxylic acid, produced by *P. aeruginosa* that are toxic to *C. elegans.*^[15]^ 1-HP is thought to act in *C. elegans* by causing α-synuclein and polyglutamine-induced protein misfolding and exacerbating α-synuclein-induced dopaminergic neurodegeneration.^[16]^ *C. elegans* modifies 1-HP by adding one, two, or three glucose moieties, with phosphorylation also observed in the endo-metabolome.^[17]^ In this study, we sought to more thoroughly characterize the metabolized 1-HP derivatives produced by *C. elegans* and to conduct preliminary experiments on the UGT gene family, which is likely involved in at least some of these modifications.

Uridine 5’-diphospho-glycosyltransferases (UGTs) are a family of enzymes critical for the homeostatic regulation of endogenous metabolites and xenobiotic detoxification in several organisms, including humans and *C. elegans*.^[18, 19]^ UGTs are the primary protein family responsible for adding glucose moieties during phase-II xenobiotic detoxification in *C. elegans.*^[9]^ Loss or modification of UGTs has been implicated in drug hypersensitivity.^[20, 21]^ They have also been shown to be upregulated upon exposure to metals, pathogenic toxins, anthelmintics, and other small molecules.^[21–26]^

## RESULTS AND DISCUSSION

### Quantitation of 1-HP Derivatization in ugt mutants

In this study, we tested available *ugt* mutants for their involvement in 1-HP modification and susceptibility (Table 1, Supplemental Table 1). We analyzed the phylogeny of these *ugt* genes and found that they covered most of the clades in the UGT family ^[31]^(Supplemental Figure 1). We then adapted a plate-based mortality screen from our previous work to discover strains with modified sensitivity to 1-HP exposure at the LD50 (179 μM) concentration (Figure 1A). ^[32]^ All strains were paired with N2 and PD1074 replicates as reference controls. We found that all strains had higher mortality when exposed to 179 μM 1-HP than the bacteria control. The solvent, DMSO, did not affect worm mortality (Supplementary Figure 2B).

**Figure 1.**
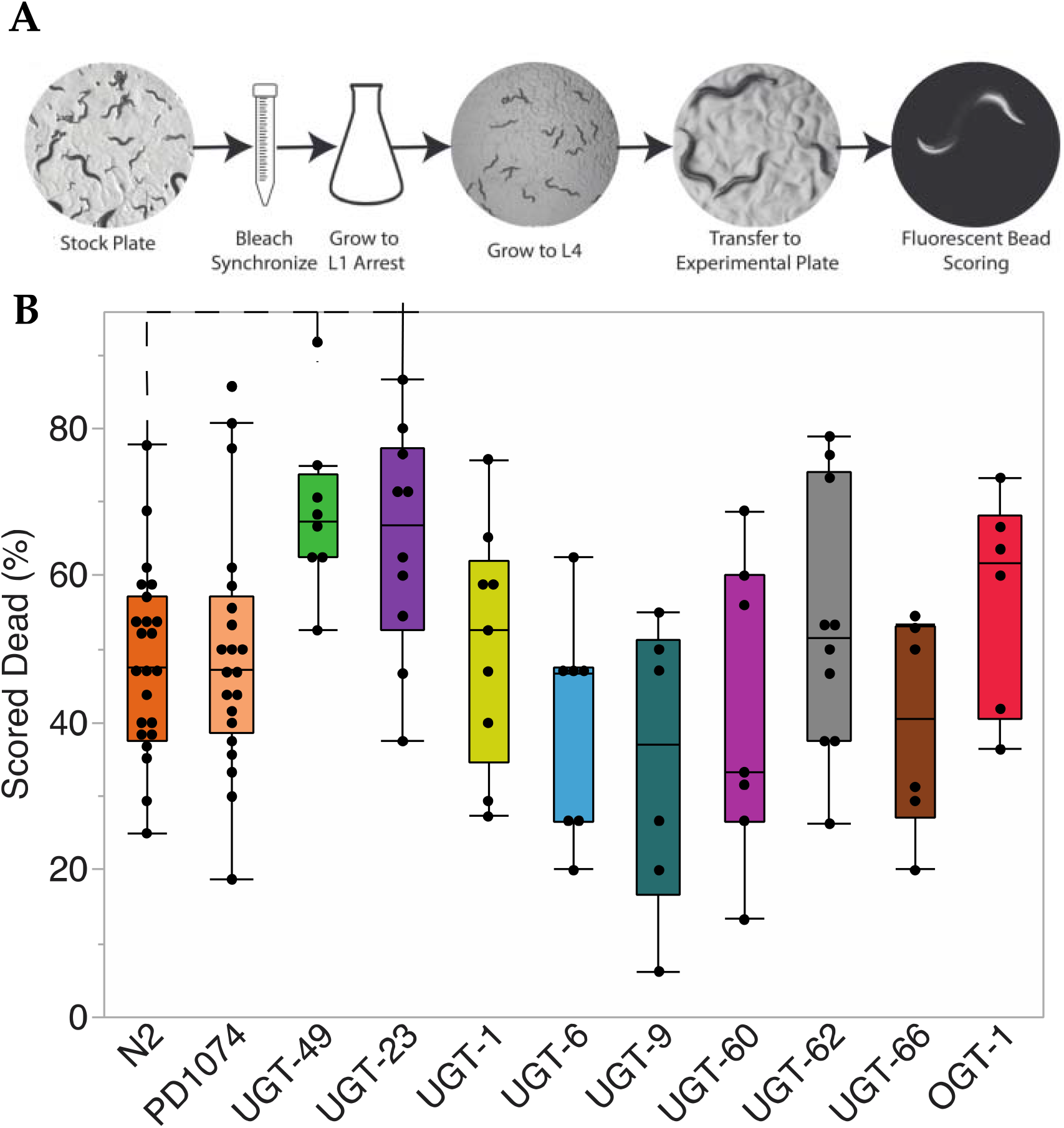
Plate-based screen for susceptibility to 1-HP. (A) Schematic describing the method for the plate-based assay. Worms were incubated for seven hours with at least six replicates per strain. (B) Box and whiskers plot with quartiles showing mortality of various strains at 179 mM 1-HP. Dashed lines indicate significantly increased mortality compared to N2 with an α of 0.1 after a Wilcoxon Pairwise comparison followed by a Benjamini-Hochberg Correction.

**Table 1.**
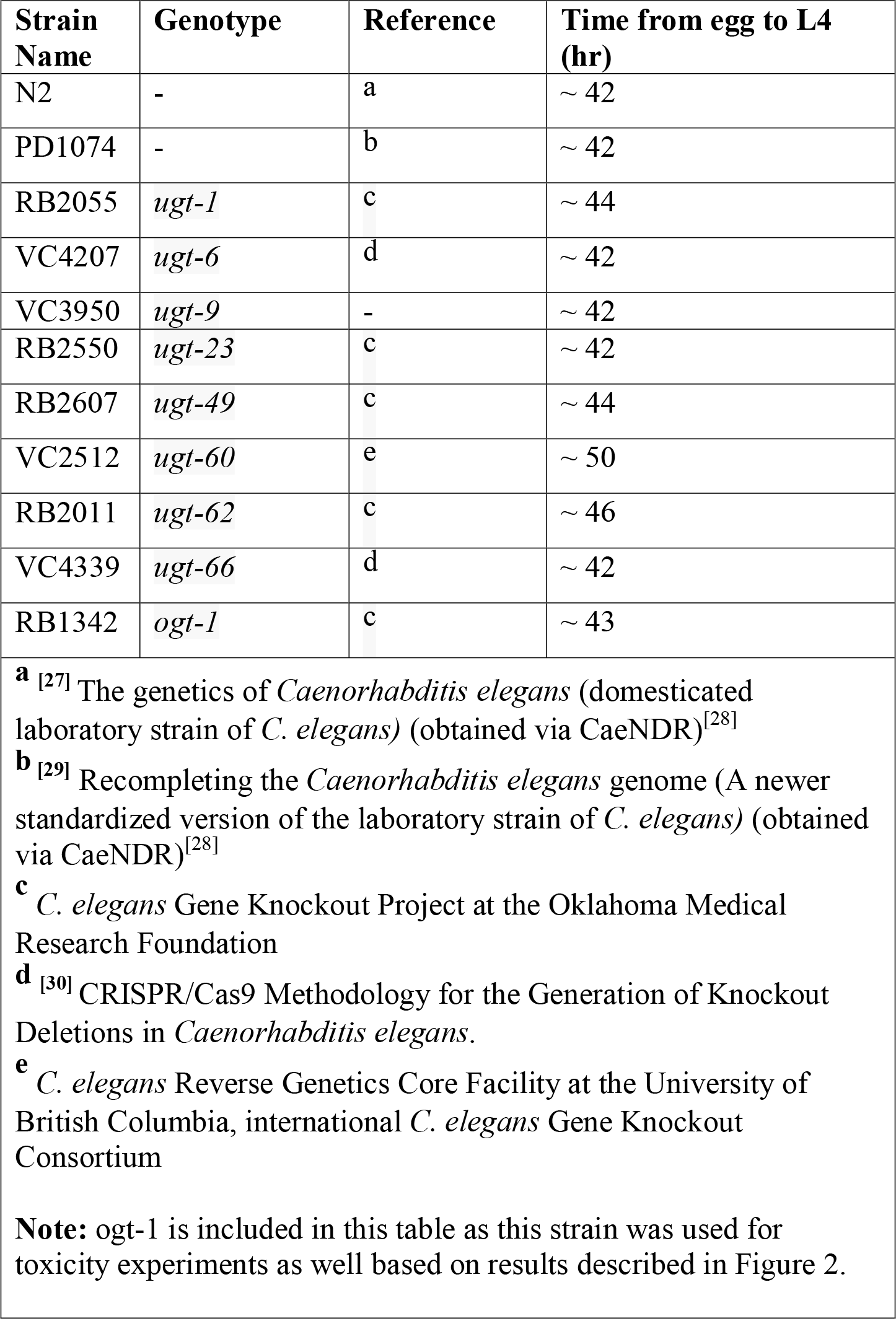
Information on strains used in this study. All strains were obtained from Caenorhabditis Genetics Center (CGC).

We tested 11 different strains for 1-HP exposure. Of these strains, N2 and PD1074 had already been tested in previous studies ^[17, 32]^. In our assay, N2 worms had a mean mortality of 48.2% with a standard error of 2.6 (n=23), while PD1074 had a mean of 49.5% with a standard error of 2.7 (n=21). We performed a one-way ANOVA in both cases, suggesting that the mortality by 1-HP exposure was significantly greater than in the controls (P < 0.0001) (Supplemental Figure 2A). In the other strains we tested (Table 1, Supplementary Table 1), we found that all the strains except *ugt-23* and *ugt-49* had a mortality percentage of between 35 and 54% and significant mortality compared to the controls (P < 0.01) (Supplemental Figure 2A). The strain *ugt-23* had a mortality of 64.7 percent with a standard error of 3.7 (n=10), and *ugt-49* had a mortality of 68.7 percent and a standard error of 3.8 (n=8). Both strains had significant mortality compared to controls (P < 0.0001). Finally, we performed a non-parametric Wilcoxon pairwise analysis for 1-HP mortality between all the strains, followed by a Benjamini-Hochberg Correction. We found that both the *ugt-23* and *ugt-49* mutants had significantly increased mortality compared to N2 (P < 0.1) (Figure 1B). This suggests that these genes have a role in the glycosylation of 1-HP.

### Isolation of glycosylated 1-HP derivatives

We then explored whether *ugt-23* and *ugt-49* produced the same 1-HP glycosylated products identified previously in N2.^[17]^ We exposed larval stage 4 (L4) worms in large-scale liquid culture for 24 hrs, followed by HPLC-UV analysis of the worm media. We found that a 22.3 μM concentration allowed many worms to survive 24 hours to accumulate sufficient modified 1-HP. Using HPLC, we observed four unique peaks in all strains (Figure 2B). The peaks were isolated using semi-preparative C-18 reverse phase HPLC and then analyzed by NMR and LC-MS/MS (Figure 2A, Supplemental Figures 2, 3, 4, 5). We identified compounds (**2**) and (**3**), which had also been identified in prior literature (Figure 2A, Supplemental Figure 2, 3).^[17]^ However, we also identified two branch-chained trisaccharides, one with three glucose moieties (**4**) and one with two glucose and an N-acetylglucosamine (GlcNAc) (**5**) (Figure 2A).

**Figure 2.**
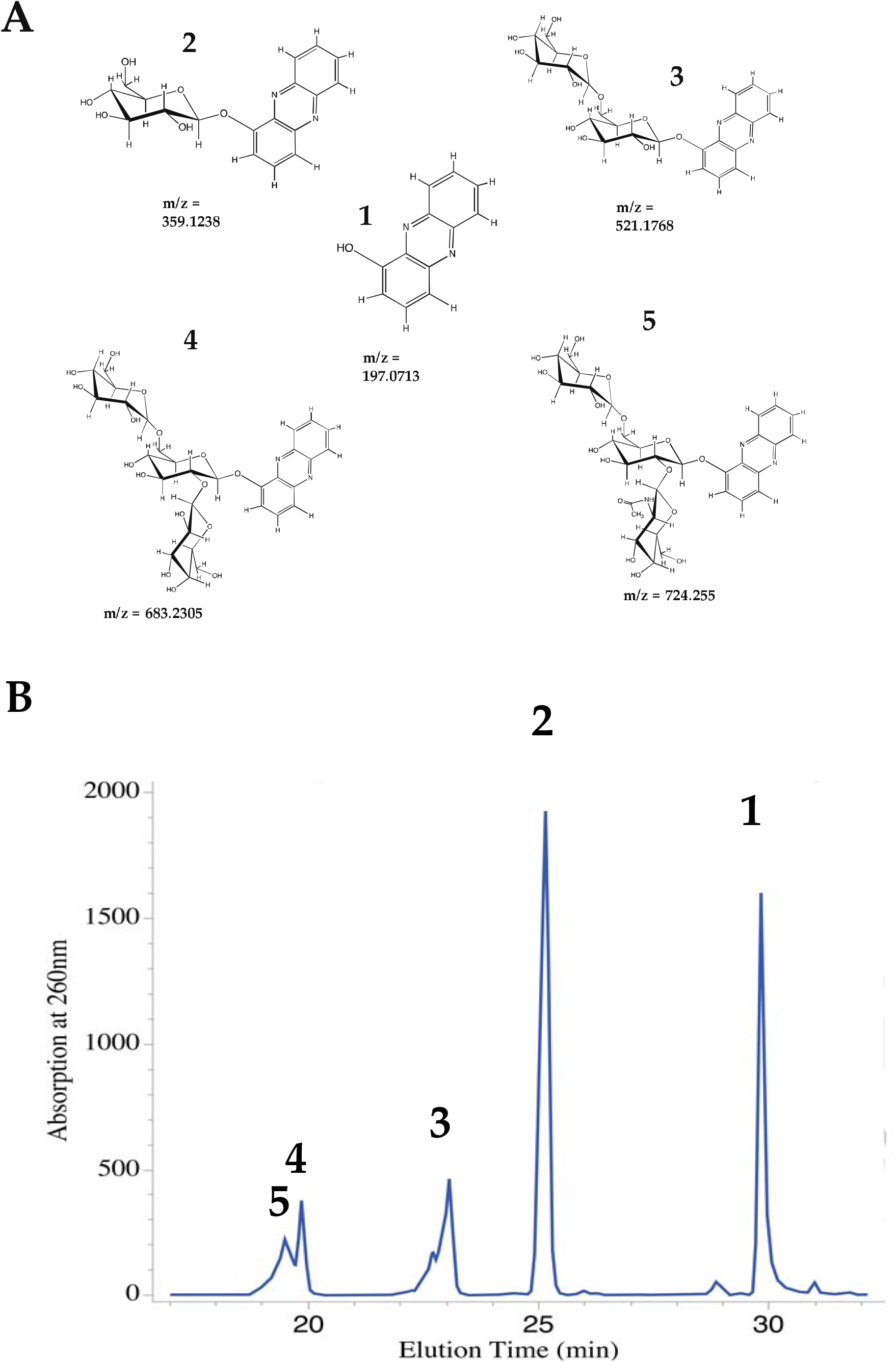
Compounds detected during the analysis of culture supernatant from PD1074 exposed to 1-HP. (A) 1-HP and its glycosylated derivatives with their corresponding m/z values. (**1**) 1-HP, (**2**) b-D-glucopyranosyl-phenazine (**3**) b-D-glucopyranosyl (1-6)-b-D-glucopyranosyl-phenazine, (**4**) b-D-glucopyranosyl (1-6)-[b-D-glucopyranosyl (1-2)]-b-D-glucopyranosyl-phenazine, and (**5)** b-D-glucopyranosyl (1-6)-[b-D-N-acetylglucosamine-pyranose (1-2)]-b-D-glucopyranosyl-phenazine. The m/z values were obtained from high-resolution MS data acquired using positive-ion electrospray ionization (ESI) (Supplemental Table 2). (B) Representative UV chromatogram of PD1074 exposed to 22.3 mM 1-HP for 24 hrs. in an S-basal medium with 2% *E. coli*. Each peak corresponds to either 1-HP or one of its glycosylated derivatives. The peak at 30 min. is (**1**), the peak at 25 min is (**2**), the peak at 23 min is (**3**), the peak at 19.8 min is (**4**), and the peak at 19.4 min is (**5**).

### Mass Spectrometry Analysis

Compounds (**4)** and (**5**) were obtained from fractionating 1-HP derivatives. Molecular compositions were determined by accurate mass measurement and tentative molecular structures were established using tandem mass spectrometry (MS/MS) (Figure 3A, 3B). The observed *m/z* of compound (**4)** was 683.2305 (theoretical value 683.2294, mass measurement error = 1.61 ppm) (Figure 2A, Supplemental Table 2). The molecular formula was established as C_30_H_38_N_2_O_16._ The observed *m/z* of compound (**5)** was 724.255 [M + H^+^] (theoretical value 724.2559, mass measurement error = 1.24 ppm (Figure 2A, Supplemental Table 2), and the molecular formula was assigned as C_32_H_41_N_3_O_16_.

**Figure 3.**
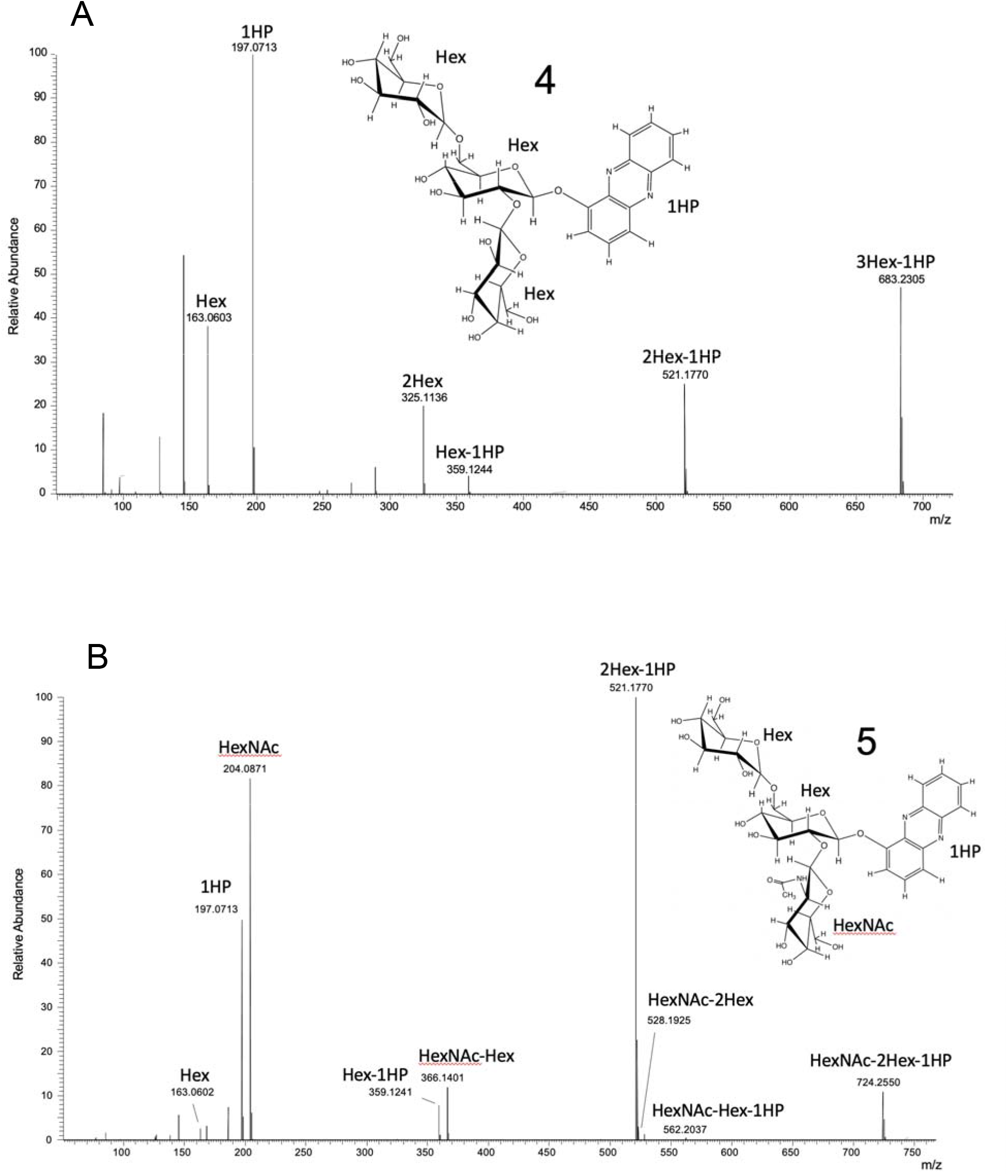
Tandem mass spectrometry (MS/MS) data for Compounds (**4**) and (**5**). Tentative structural insets are based on NMR results discussed in detail below. (A) MS/MS Spectra for compound (**4**) with fragment ions corresponding to the loss of three hexose sugars. (B) MS/MS Spectra for compound (**5**) with fragment ions corresponding to the loss of two hexose and an N-Acetyl hexose sugar.

HCD fragmentation of these compounds resulted in the generation of largely B-type glycosidic bond cleavages that enable the partial assignment of molecular structure. MS/MS of compound (**4)** produced multiple fragment ions that support the modification of 1-HP by three hexose sugar units with the primary fragments annotated in Figure 3A. The sequential neutral loss of three hexose sugars due to glycosidic bond cleavage from the isolated molecular ion was observed, leading to a fragment identified as 1-HP (observed m/z 197.0713). Additional fragment ions corresponding to hexose units are also observed. The MS/MS of Compound (**5**) produced multiple fragment ions shown in Figure 3B that support the modification of 1-HP by two hexose sugars and one N-acetyl hexose sugar. Similar to the fragmentation observed in compound (**5**), the sequential neutral losses of sugar residues via glycosidic bond cleavage from the modified 1-HP molecular ion are also observed, with unique ions now present due to the inclusion of N-Acetyl moiety. A table of assignment fragments is provided as Supplemental Table 3.

### Nuclear Magnetic Resonance Analysis

The ^1^H NMR spectrum of compound (**4**) (Figure 4A) contained signals corresponding to a phenazine, a sugar region with three anomeric protons, including two (H1’ and H1’’) that are unusually downfield for beta-anomers but consistent with being close to the aromatic phenazine. Analysis of data from 2D TOCSY and COSY experiments (Supplemental Figure 5.1) indicates that the three protons identified in the ^1^H spectrum belong to three glucose moieties. The glucosyl attached to the phenazine and the glucosyl linked at C6 have chemical shifts very similar to those of the gentiobiose-phenazine described in Stupp et al.^[17]^ The third glucosyl attached to C2 has very unusual shifts, consistent with proximity to the phenazine, but the coupling patterns match the glucose configuration. Figure 4B illustrates the linkage position of compound (4), determined using 1D and 2D ROESY data. The bottom panel shows a region from the TOCSY spectrum with H1’ along the horizontal at 5.77 ppm coupled to H2’-H6’; assignments are annotated in the top panel. This glucose is linked to the phenazine at the position shown in Figure 4A (see also supplemental Figure 5.2). The middle panel shows a region from the ROESY spectrum with an NOE cross peak from H1’’ at 5.1 ppm to H2’, establishing the H1’’ – H2 linkage. Similarly, the H1’’’ proton at 4.469 ppm shows cross-peaks linking it to H6’ of the same phenazine-linked glucose.

**Figure 4.**
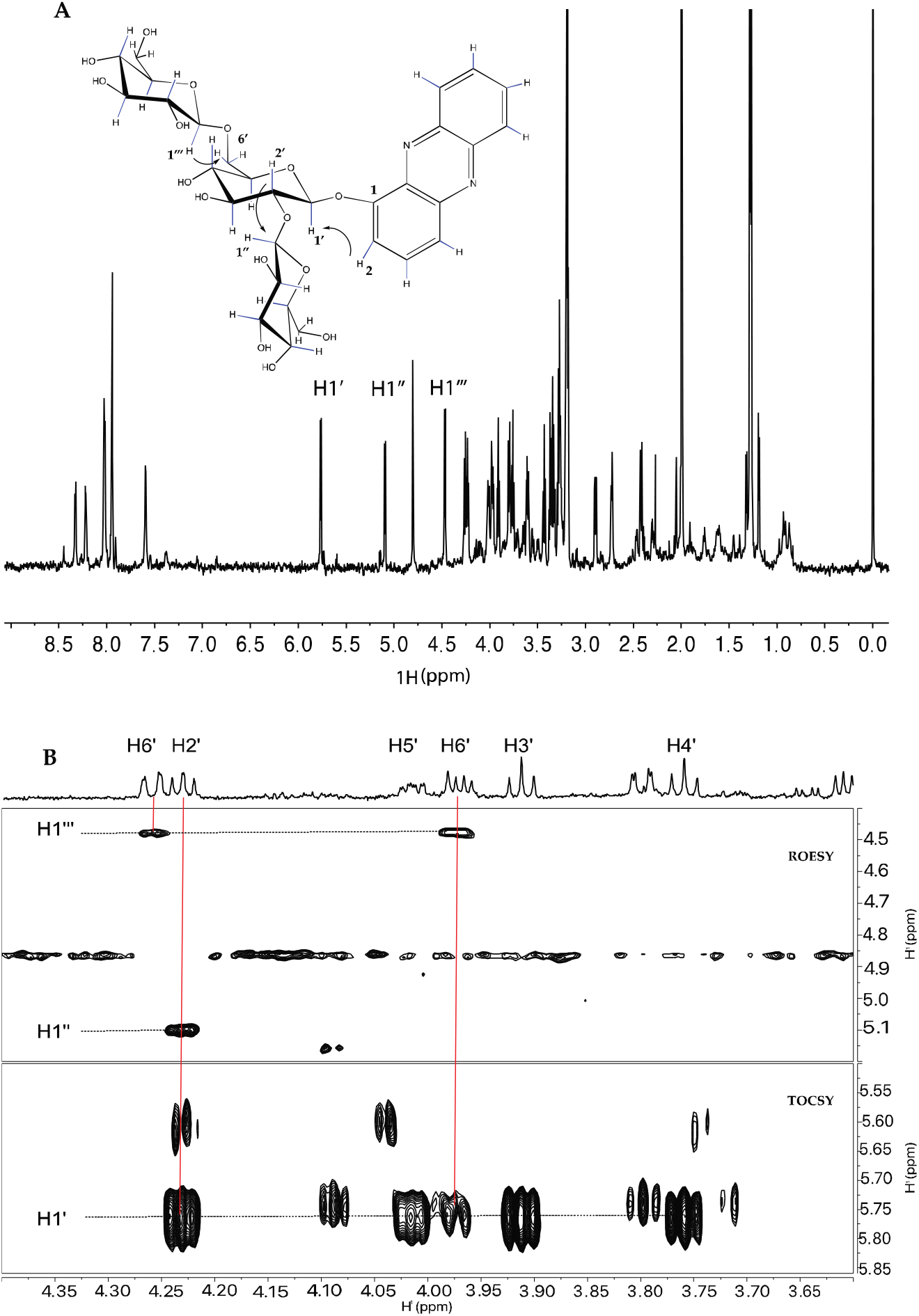
NMR spectra for trisaccharide compound (**4**): (A) Structure of compound (**4)** and 1D proton spectrum with b-glucosyl anomeric protons annotated. (B) The top panel shows a 1D proton of the glucosyl residue attached to the phenazine. The bottom panel is a region of a 2D TOCSY showing the protons coupled to H1’. The middle panel is a region of a 2D ROESY showing NOEs between H1’’ and H1’’’ and the respective protons in the linkage positions.

The ^1^H NMR spectrum of the minor compound (**5**) (Figure 5A) similarly displayed signals corresponding to the phenazine and three sugar residues. However, a difference in chemical shift of H2’’ (Figure 5B) suggested a β-NAcetyl-2-deoxy-glucosamine residue instead of a glucosyl residue (CASPER database),^[33]^ consistent with the MS/MS data (Figure 3B). A combination of 1D and 2D TOCSY and 1D ROESY data (Supplemental Figure 6) confirmed that the three residues were the correct configuration and the linkages were the same as in compound (**4).**

**Figure 5.**
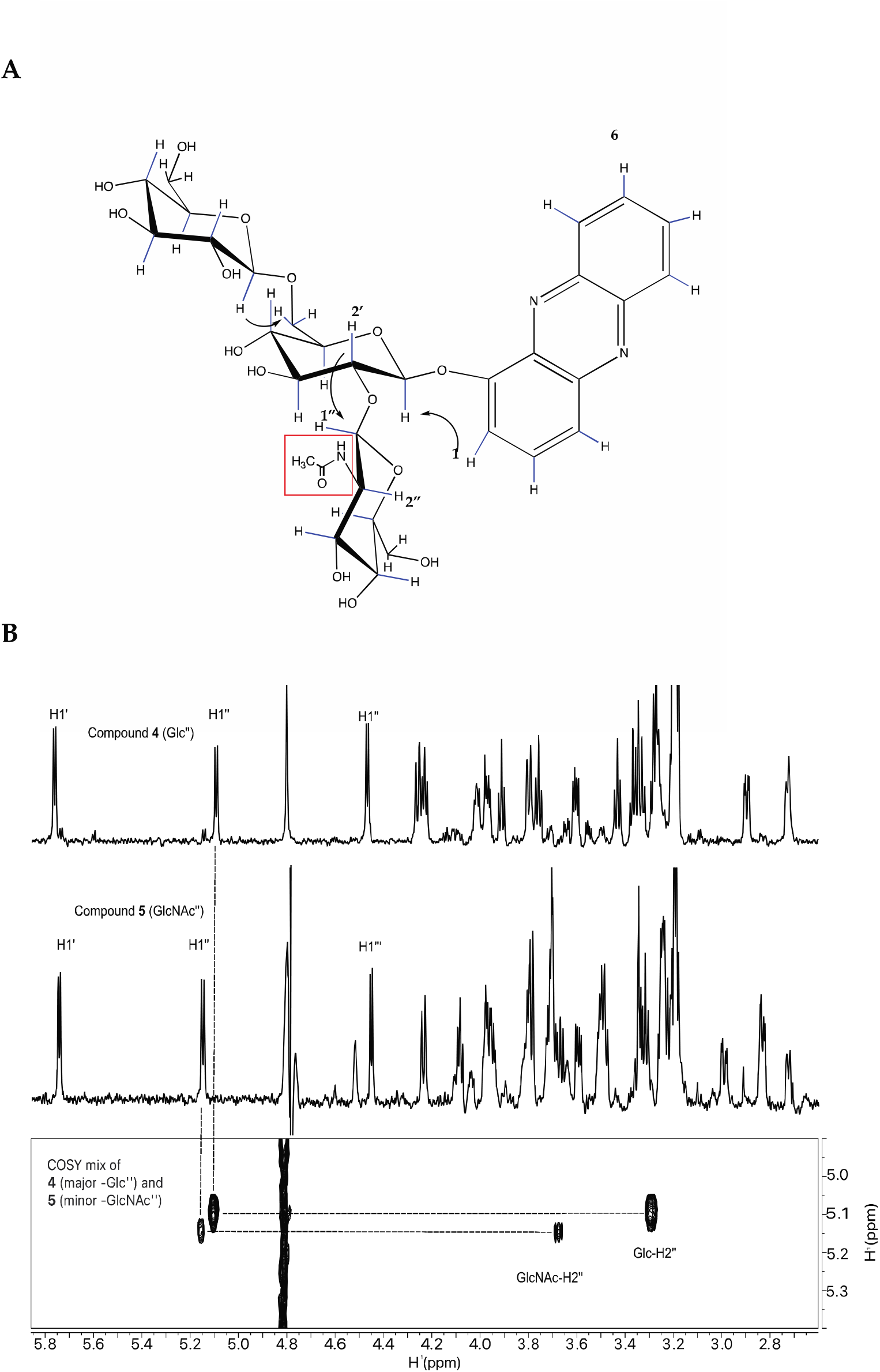
Proton spectrum supporting the NAcetyl-glucosamine containing compound (**5)**: (A) Structure of compound **5**. (B) The Middle panel shows the 1D proton of 5 and the three β-anomeric protons like those in compound (**4)** (top panel). The bottom panel is a region from a 2D COSY of the mixture of compound (**4)** (primary compound) and compound (**5)** (minor compound), indicating the chemical shift positions of H2’’.

Because (**5**) contained a GlcNAc, we evaluated the GlcNAc transferase *ogt-1* with the same assays described above. *ogt-1* has previously been shown to modulate the immune response in *C. elegans* for *S. aureus* infection but not *P. aeruginosa* infection.^[34]^ Consistent with those findings, the *ogt-1* knockout had no statistically significant difference in susceptibility to 1-HP compared with N2 and PD1074 (Figure 1B). LC-MS analysis of worm media conditioned by the *ogt-1* knockout mutant challenged with 1-HP showed that (**5**) was still produced (Supplemental Figure 7).

### Quantitation of 1-HP Derivatization in ugt knockout mutants

We then quantified the HPLC-UV data to observe if there was a reduction in the amounts of 1-HP derivatives for the *ugt-23* and *ugt-49* knockout mutants. We normalized the data to the sum of all the 1-HP-related compounds for each replicate. This ensured that the ratio obtained was independent of any variation due to the amount of 1-HP the worms were exposed to or the number of worms per replicate. We found that the *ugt-23* knockout mutant produced decreased amounts of both trisaccharide sugars (**4**) and (**5**), while the *ugt-49* knockout mutant had reduced amounts of compounds (**3**), (**4**), and (**5**) (Figure 5).

**Figure 5.**
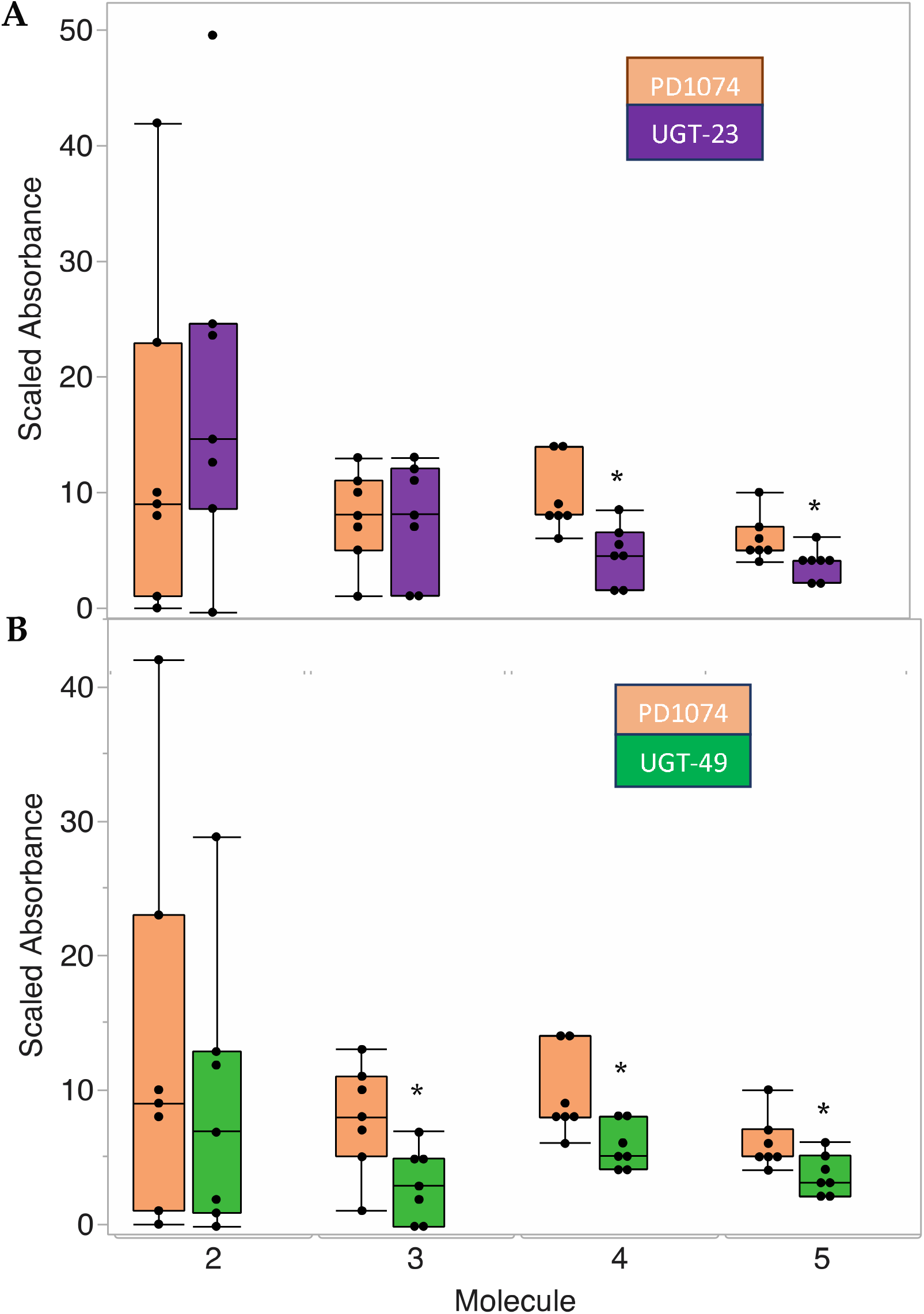
Box and whiskers plot showing the relative amounts of 1-HP and its derivatives after a 24-hour incubation at 22.3 mM 1-HP with 2% *E. coli* based on UV absorbance data (n=7). All replicates were paired, and data were normalized by dividing the absorbance for each compound at 260 nm by the sum of the absorbances of 1-HP and all its derivatives for each run (abs *x*/[abs *z* + abs *y* + abs *x* + abs *w* + abs *v*]). * Indicates significant difference in relative amounts of compound compared to the relative amount of the same compound in PD1074 after Wilcoxon pairwise analysis (α = 0.05). (A) Relative amounts of glycosylated 1-HP derivatives for the *ugt-23* mutant compared to PD1074. Compounds **4** and **5** are reduced in this strain. (B) Relative amounts of glycosylated 1-HP derivatives for the *ugt-49* mutant compared to PD1074. Compounds **3**, **4**, and **5** are reduced in this strain.

These results show the involvement of *ugt* genes in 1-HP detoxification, suggesting that they have broad specificity and that multiple *ugt* genes are involved in detoxifying a xenobiotic in *C. elegans*. Prior research has implicated multiple *ugt* genes responsible for the glycosylation of other small molecule toxins such as indole.^[35]^ The workflow outlined in this study can be used to test the role of *ugt* genes in modifying other small molecule xenobiotics. Future studies could validate whether broad specificity is seen in response to xenobiotics or whether this is a phenomenon specific to 1-HP.

Furthermore, our results implicate the addition of GlcNAc in detoxification in *C. elegans,* a result which, to our knowledge, has yet to be previously observed. Our data also suggest that genes other than *ogt-1* are responsible for adding GlcNAc in the 1-HP glycosylation pathway. This might be due to the broad specificity of *ugt* genes or that GlcNAc serves a particular purpose in detoxification. Using this workflow, it would be interesting to see if GlcNAc-modified products are also observed for other xenobiotics.

## METHODS

### Mortality Assay

All 11 strains of *C. elegans* were grown and maintained on 10 cm NGM agar plates seeded with an LB-cultured OP50 strain of *E. coli* at 22 C. Knockout mutants were paired with an N2, and PD1074 replicates for each strain. 10 cm NGM plates with *C. elegans* were bleached and grown to L1 arrest, and then L1 arrested worms were transferred to new 10 cm plates and allowed to grow to L4. Upon reaching L4, ∼ 15 worms were assigned either to control 6 cm plates or 6 cm plates with NGM and 179 μM 1-HP, the LD50 value of PD1074.^[22]^ Worms were incubated on experimental plates for 7 hours. After 7 hours, fluorescent beads were added to the worms, and the uptake of these beads was used as a marker to differentiate between alive and dead worms.^[36]^

### Lifespan Timing Assay

Worms were observed to determine how long they took to go from egg to L4 in two different ways. The time to L4 for N2, PD1074, and the *ugt-1, ugt-23, ugt-49, ugt-60,* and *ugt-62* knockout mutants was measured by initially spot bleaching a single adult and following a single egg, observing them until they reached L4. The time to L4 for the *ugt-6, ugt-9, ugt-66, and ogt-1* knockout mutants was measured by bleach synchronizing a plate of worms and placing the resulting eggs on a 10 cm plate. The plates were observed every four to eight hr. until most of the population on the plate could reliably be identified as L4.

### Large Scale Growth of *C. elegans*

Worms were grown on large-scale culture plates (LSCPs) to generate worms for subsequent experiments. LSCPs were poured according to previously described protocols.^[37]^ Poured plates were seeded with the HTS115 strain of *E. coli* prepared in K-media at a concentration of 0.5 g mL^-1^ bacteria generated according to the IBAT method.^[38]^ Worms were chunked onto the LSCPs and then grown for seven to ten days, depending on the strain, before being washed with M9 for subsequent experiments. After washing, worms were bleach synchronized and then grown to L1 arrest in M9. Upon reaching L1 arrest, they were transferred to an S-basal medium (∼30,000 worms mL^-1^) and incubated with 2 % *E. coli* OP50 until they reached L4. After the worms had reached L4, they were incubated with either 1.1 % DMSO or 22.3 μM 1-HP. Worms were incubated for 24 hours and then centrifuged. The supernatant was collected for subsequent experiments.

### Glucoside Collection and Analysis (HPLC-UV)

After the supernatant was separated from the worms, it was centrifuged again at 20,800 RCF for 10 minutes to separate the bacteria from the supernatant. The resultant volume was lyophilized and extracted in an appropriate volume of methanol (200-600 μL, depending on the starting volume of the supernatant). It was then centrifuged at 20,800 RCF for 30 minutes. Following centrifugation, the supernatant was concentrated to ∼100 μL with 90 μL injected into the HPLC-UV and 10 μL separated for LC-MS.

The supernatant was analyzed on an Agilent 1200 Series HPLC system with a diode array collector, and fractions were collected manually upon observing a peak. Absorbance was measured at 260nm. For worm media separation, 5% methanol (A) and 95 % 5mM Phosphate buffer pH 7.2 (B) were held isocratic for four minutes, increasing to 95 % A and 5 % B over 30 minutes, and then held for five minutes, followed by a re-equilibration of the column. The separation was carried out at a flow rate of 2 mL min^-1^ in an Agilent SB C-18 column (9.4 mm x 250 mm, 5μM).

After initial fractionation, further separation of the fraction containing compounds 4 and 5 was carried out. For that separation, 5% methanol (A) and 95% five mM Phosphate buffer pH 7.2 (B) were held isocratic for four minutes, increasing to 50 % A and 50 % B over 17 minutes. The gradient was slowed, and the ratio increased to 67% A and 33% B by 28 minutes before ramping it up to 95% A and 5% B by 30 minutes and then holding constant for five minutes. A re-equilibration of the column followed this. The column used for this separation was the same as for the initial worm media separation.

### Glucoside Analysis (LC-MS/MS)

Samples aliquoted during glucoside collection were analyzed using a Thermo Fisher Scientific Q Exactive HF Orbitrap mass spectrometry coupled to a Vanquish UPLC with inline UV detection. Chromatographic separation was performed with an Agilent ZORBAX Eclipse XDB-C18 column (2.1×150mm, 1.8 micron) over 30 minutes, starting with 95 % H_2_O (A) and 5% methanol (B) held isocratic for 2.5 minutes, then increased to 70% B by 22 minutes and 100% B at 22.5 minutes and held for 4 minutes before re-equilibration at 5% B for 3 minutes prior the next injection. The sample queue was randomized with injection blanks included to monitor for sample carryover. All samples were analyzed by positive mode electrospray ionization (ESI). Full MS scans were performed at a specified resolution of 30,000 (m/z 200) from 150 to 2,000 m/z with an AGC target of 3e6 and a maximum IT of 200 ms. Corresponding UV traces were collected at 260 nm.

Target compounds were isolated with a 4.0 m/z quadrupole window to perform structural elucidation by higher-energy collisional dissociation (HCD). A normalized collision energy (NCE) of 15 V was applied, and fragment ions were detected with a specified resolution of 15,000, AGC target of 2e5, and a maximum of IT 100 ms. MS data were analyzed with Thermo Qual Browser and manually interpreted.

### Glucoside Analysis (NMR)

Pooled fractions were dried, resuspended in 60 μL D_2_O with 0.15 mM DSS as an internal standard, dried with a speed vac, and resuspended twice in 60 μL D_2_O to perform buffer exchange to remove excess H_2_O before being transferred into 1.7 mm NMR tubes. 1D ^1^H, 2D COSY, 2D TOCSY, selective 1D TOCSY, and selective 1D ROESY spectra were collected where appropriate on a Bruker 800 MHz NEO spectrometer using a 1.7 mm cryoprobe. Spectra were processed and analyzed with MestReNova 14.1.2 (Mestrelab Research).

### Statistical Analysis

Analysis was performed using JMP^®^, a publicly available statistical software. A Wilcoxon test, followed by a Wilcoxon pairwise analysis and a Benjamini-Hochberg correction, was performed for the mortality assays to determine the significance between strains. Tukey’s HSD test was performed to determine the significance within each strain for 1-HP exposure. A Wilcoxon test followed by Wilcoxon pairwise analysis was performed on the scaled absorbance data.

## Notes

The Authors declare no competing financial interests.

## ACKNOWLEDGEMENTS

We thank Olatomiwa Bifarin (Georgia Tech) for helpful discussions on designing worm toxicity experiments. We thank Gonçalo Gouveia (NIST), Amanda Shaver (Northwestern University), and Pam Kirby (University of Georgia) for help and discussions on large-scale worm growth protocols. We thank Laura Morris and Ricardo Borges (University of Georgia) for their help using the HPLC instrument and for valuable data analysis discussions. This work was supported by NIH 1U2CES030167 and the Georgia Research Alliance.

