## Supplemental Figures for "Role of *ugt* genes in detoxification and glycosylation of 1-hydroxy phenazine (1-HP) in *Caenorhabditis elegans*"

**Supplemental Table 1.** Detailed information on strains used in this study, including strain name, genotype affected, gene sequence, deletion strategy, reference article where strain is first mentioned, and time each strain takes to grow to L4.

| **Strain Name** | **Genotype** | **Gene Sequence** | **Description** | **Reference Article** | **Hours from egg to L4** |
| --- | --- | --- | --- | --- | --- |
| N2 | - | - | - | (Brenner, 1974) | ~ 42 hours |
| PD1074 | - | - | - | (Yoshimura et al., 2019) | ~ 42 hours |
| RB2055 | ugt-1(ok2718) V. | AC3.7 | Homozygous. Outer Left Sequence: AGCCATGAGGACAAAGTTCG. Outer Right Sequence: TGTTGAAAATGCTTTGCCAG. Inner Left Sequence: TTGCTCTTCCATGTCTCGAA. Inner Right Sequence: TCGTCTAGATTCCGCCATTT. Inner Primer PCR Length: 1255 bp. Deletion Size: 787 bp. Deletion left flank: TTGCATTTCCAACACCATCACTTCCAAAAC. Deletion right flank: TTTGTTCAGTACTACATGTTAGATGCTTTT. | C. *elegans* Gene Knockout Project at the Oklahoma Medical Research Foundation,  International C. *elegans* Gene Knockout Consortium | ~ 44 hours |
| VC4207 | ugt-6(gk5292[loxP + myo-2p::GFP::unc-54 3' UTR + rps-27p::neoR::unc-54 3' UTR + loxP]) V. | ZC455.4 | Homozygous viable. Deletion of 1469 bp with Calarco/Colaiacovo selection cassette conferring myo-2 GFP and G418 resistance inserted at break. Left flanking sequence: GCGTGTTTTACCATCAGATCAGTTGGCGTG ; Right flanking sequence: GATGGAATATGCTGCCTTCCAAGTTCATCA. | (Au et al., 2019) | ~ 42 hours |
| VC3950 | ugt-9(gk5024[loxP + myo-2p::GFP::unc-54 3' UTR + rps-27p::neoR::unc-54 3' UTR + loxP]) V. | T19H12.1 | Homozygous viable. Deletion of 1266 bp with Calarco/Colaiacovo selection cassette conferring myo-2 GFP and G418 resistance inserted at break. Left flanking sequence: AAGTTTTCGGAAGGCTTTCTGTAGGGTGAA; Right flanking sequence: GGAGGTGCTGTTGCGTACGACAAATTTGAT. | - | ~ 42 hours |
| RB2550 | ugt-23(ok3541) X. | C17G1.3 | Homozygous. Outer Left Sequence: cgtgacgctttagcatttca. Outer Right Sequence: tcattgatgccgatgaagaa. Inner Left Sequence: ttgatcagcgaatattggga. Inner Right Sequence: atgcacattctcatcttgcg. Inner Primer PCR Length: 1197. Estimated Deletion Size: about 300 bp. | C. *elegans* Gene Knockout Project at the Oklahoma Medical Research Foundation,  International C. *elegans* Gene Knockout Consortium | ~ 42 hours |
| RB2607 | ugt-49(ok3633) V. | AC3.2 | Homozygous. Outer Left Sequence: cgtgtgatggtgacaagacc. Outer Right Sequence: agaacagcaacgaacacgaa. Inner Left Sequence: acgtggcattcagtgaacaa. Inner Right Sequence: ggacaaaagcaataacatcaaga. Inner Primer PCR Length: 1279. Estimated Deletion Size: about 400 bp. | C. *elegans* Gene Knockout Project at the Oklahoma Medical Research Foundation,  International C. *elegans* Gene Knockout Consortium | ~ 44 hours |
| VC2512 | ugt-60(ok3248) III/hT2 [bli-4(e937) let-?(q782) qIs48] (I;III). | C07A9.6 | Homozygous lethal deletion chromosome balanced by bli-4- and GFP-marked translocation. Heterozygotes are WT with pharyngeal GFP signal, and segregate WT GFP, arrested hT2 aneuploids, and non-GFP ok3248 homozygotes (probable early larval arrest). Homozygous hT2[bli-4 let-? qIs48] inviable. Pick WT GFP and check for correct segregation of progeny to maintain. External left primer: GAAGGTTTCGGACTTGTTGC. External right primer: CGCATCCACTTTCTTCAGGT. Internal left primer: CTGAGAGCATCGCGGATAGT. Internal right primer: TGACGCGTCTAGCTCAATTTT. Internal WT amplicon: 1354 bp. Deletion size: 525 bp. Deletion left flank: TATAGCCTCCATGTGCAATCATTAATTTCA. Deletion right flank: AACCTCGATAGAACAAATTCTCGTCAACGA. | C. *elegans* Reverse Genetics Core Facility at the University of British Columbia,  international C. *elegans* Gene Knockout Consortium | ~ 50 hours |
| RB2011 | ugt-62(ok2663) III. | M88.1 | Homozygous. Outer Left Sequence: AAACATGGTTCCCGACATTC. Outer Right Sequence: AATTCGGTGCATTTGGAAAA. Inner Left Sequence: GCAACTTTGGAATTTTTGGG. Inner Right Sequence: ATCAGATTTCCTGCGCAACT. Inner Primer PCR Length: 3024 bp. Deletion Size: 1367 bp. Deletion left flank: ACGCTAAATTGTTTTAATACATTTTAAAGT. Deletion right flank: ATGAAATATTTCTCGATTAAAGTTTCTCAG. | C. *elegans* Gene Knockout Project at the Oklahoma Medical Research Foundation,  International C. *elegans* Gene Knockout Consortium | ~ 46 hours |
| VC4339 | ugt-66(gk5422[loxP + myo-2p::GFP::unc-54 3' UTR + rps-27p::neoR::unc-54 3' UTR + loxP]) III. | C23G10.6 | Deletion of 2473 bp with Calarco/Colaiacovo selection cassette conferring myo-2::GFP and G418 resistance inserted at break. Left flanking sequence: AAAATTTCAAAATATTAAATGAAGCCGTTG; Right flanking sequence: CAGGGAGGTGTCACAATTATTTGTGTCCTG. | (Au et al., 2019) | ~ 42 hours |
| RB1342 | ogt-1(ok1474) III. | K04G7.3 | Homozygous. Outer Left Sequence: gccaaagaattgatttcgga. Outer Right Sequence: tgctcttgcaccacaaccta. Inner Left Sequence: acctgtccgagaccattctg. Inner Right Sequence: ccaacgctattgctcctctc. Inner Primer PCR Length: 2730. Estimated Deletion Size: about 1300 bp. | C. *elegans* Gene Knockout Project at the Oklahoma Medical Research Foundation,  International C. *elegans* Gene Knockout Consortium | ~ 43 hours |

**Note:** All strains were obtained from Caenorhabditis Genetics Center (CGC).

**Supplemental Table 2.** Assigned molecular compositions in high-resolution MS data for 1-HP derived C. elegans metabolites acquired using positive-ion electrospray ionization (ESI).

| **Compound** | **Ion** | **Ion Formula** | **Calculated *m/z*** | **Observed *m/z*** |
| --- | --- | --- | --- | --- |
| b-D-glucopyranosyl-phenazine | [M+H]^+^ | C_18_H_19_N_2_O_6_^+^ | 359.1237 | 359.1240 |
| b-D-glucopyranosyl (1-6)-b-D-glucopyranosyl-phenazine | [M+H]^+^ | C_24_H_29_N_2_O_11_^+^ | 521.1766 | 521.177 |
| b-D-glucopyranosyl (1-6)- [b-D-glucopyranosyl (1-2)]-b-D-glucopyranosyl-phenazine | [M+H]^+^ | C_30_H_39_N_2_O_16_^+^ | 683.2294 | 683.2305 |
| b-D-glucopyranosyl (1-6)- [b-D-N-acetylglucosamine-pyranose (1-2)]-b-D-glucopyranosyl-phenazine. | [M+H]^+^ | C_32_H_42_N_3_O_16_^+^ | 724.2560 | 724.2559 |

**Note:** Representative data is shown for N2.

**Supplemental Table 3.** Assigned fragment ions in high-resolution MS/MS data for 1-HP-derived *C.* *elegans* metabolites acquired using positive-ion electrospray ionization (ESI).

| **Compound 5** |  |  |  |
| --- | --- | --- | --- |
| Ion | Theoretical m/z | Observed m/z | Mass Measurement Error (ppm) |
| HexNAc-2Hex-1HP | 724.2559 | 724.255 | 1.24 |
| HexNAc-1Hex-1HP | 562.2031 | 562.2037 | -1.07 |
| HexNAc-2Hex | 528.1923 | 528.1925 | -0.38 |
| 2Hex-1HP | 521.1765 | 521.177 | -0.96 |
| HexNAc-Hex | 366.1395 | 366.1401 | -1.64 |
| Hex-1HP | 359.1237 | 359.1241 | -1.11 |
| HexNAc | 204.0867 | 204.0871 | -1.96 |
| 1HP | 197.0709 | 197.0713 | -2.03 |
| 1Hex | 163.0601 | 163.0602 | -0.61 |
| **Compound 4** |  |  |  |
| Ion | Theoretical m/z | Observed m/z | Mass Measurement Error (ppm) |
| 3Hex-1HP | 683.2293 | 683.2305 | -1.76 |
| 2Hex-1HP | 521.1765 | 521.177 | -0.96 |
| 1Hex-1HP | 359.1237 | 359.1136 | 28.12 |
| 2Hex | 325.1129 | 325.1136 | -2.15 |
| 1HP | 197.0709 | 197.0713 | -2.03 |
| 1Hex | 163.0601 | 163.0603 | -1.23 |
| **Compound 3** |  |  |  |
| Ion | Theoretical m/z | Observed m/z | Mass Measurement Error (ppm) |
| 2Hex-1HP | 521.1765 | 521.1768 | -0.58 |
| Hex-1HP | 359.1237 | 359.1245 | -2.23 |
| 1HP | 197.0709 | 197.0713 | -2.03 |
| 1Hex | 163.0601 | 163.0603 | -1.23 |
| **Compound 2** |  |  |  |
| Ion | Theoretical m/z | Observed m/z | Mass Measurement Error (ppm) |
| Hex-1HP | 359.1237 | 359.1238 | -0.28 |
| 1HP | 197.0709 | 197.0716 | -3.55 |

**Note:** Representative data is shown for N2.

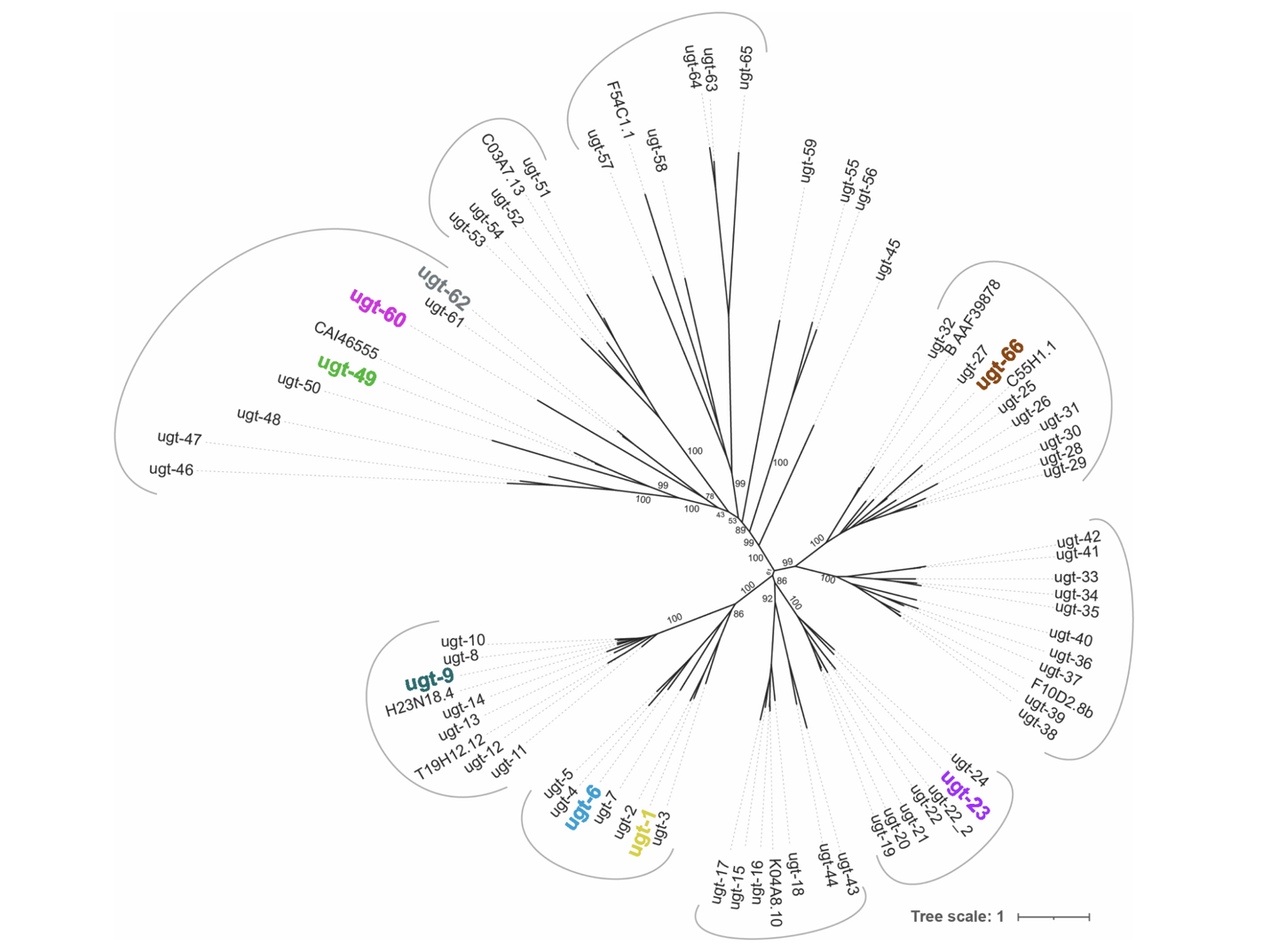

**Supplemental Figure 1.** Phylogenetic tree highlighting *ugt* genes used in this study^[1]^.

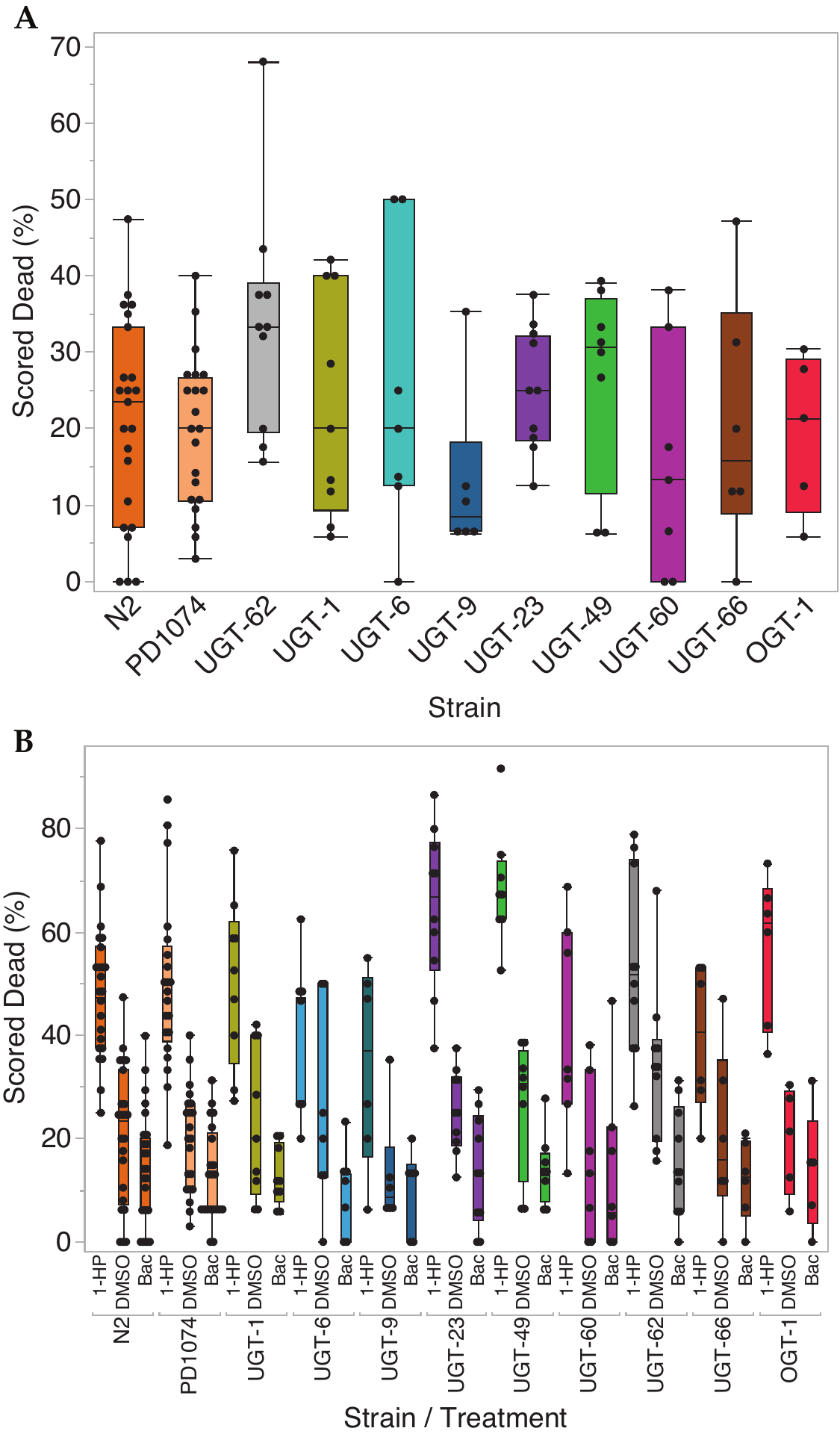

**Supplemental Figure 2.** Percentage of worms scored in plate-based mortality screen after 7 h incubation. (A) Box and whiskers plot showing mortality of various strains to 1.1% DMSO for 7h (n >/= 6). (B) Box and whiskers plot showing combined data of exposure to 179μM 1-HP, 1.1% DMSO and bacteria control showing increased mortality upon 1-HP exposure for each strain compared to controls after Tukey’s HSD test (α = 0.05).

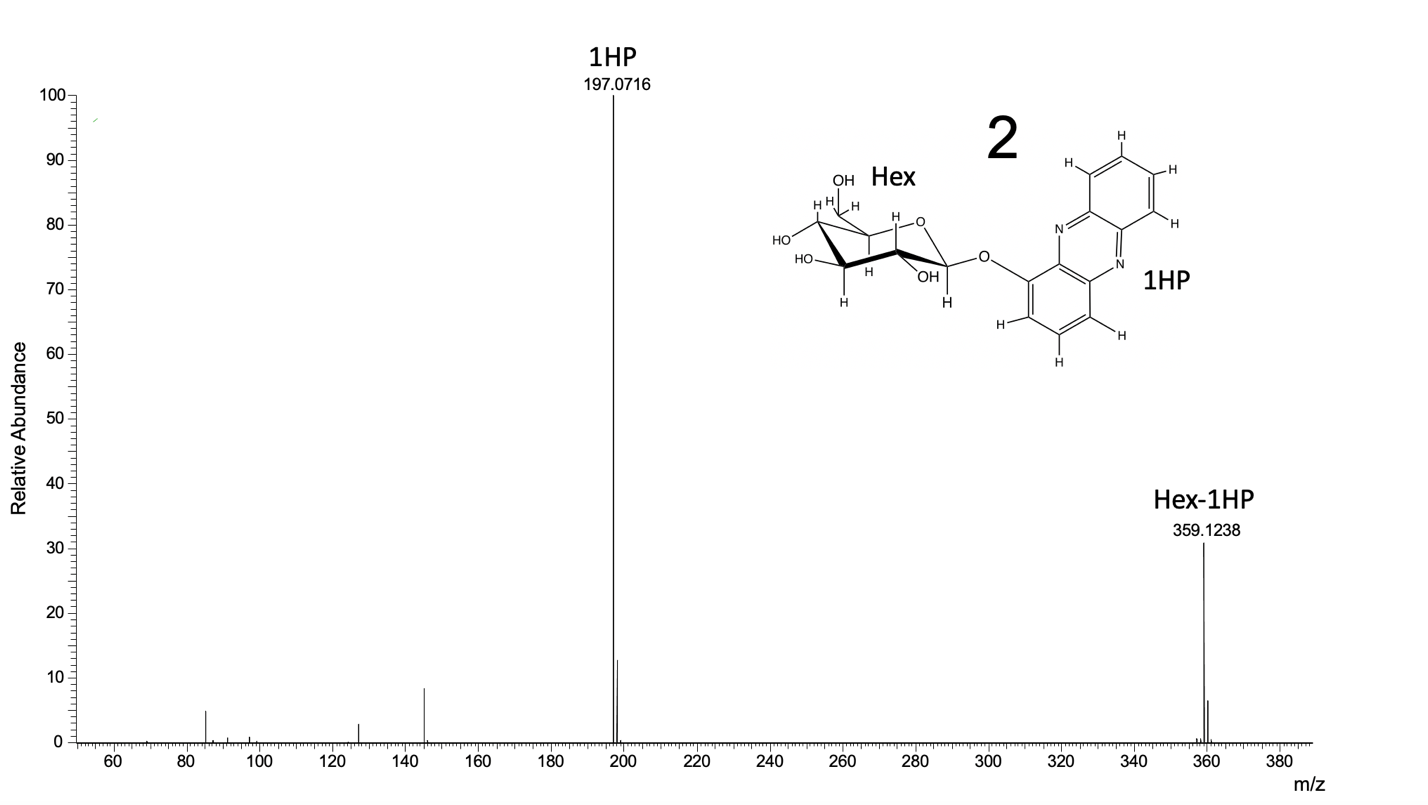

**Supplemental Figure 3.** MS/MS data for (2).

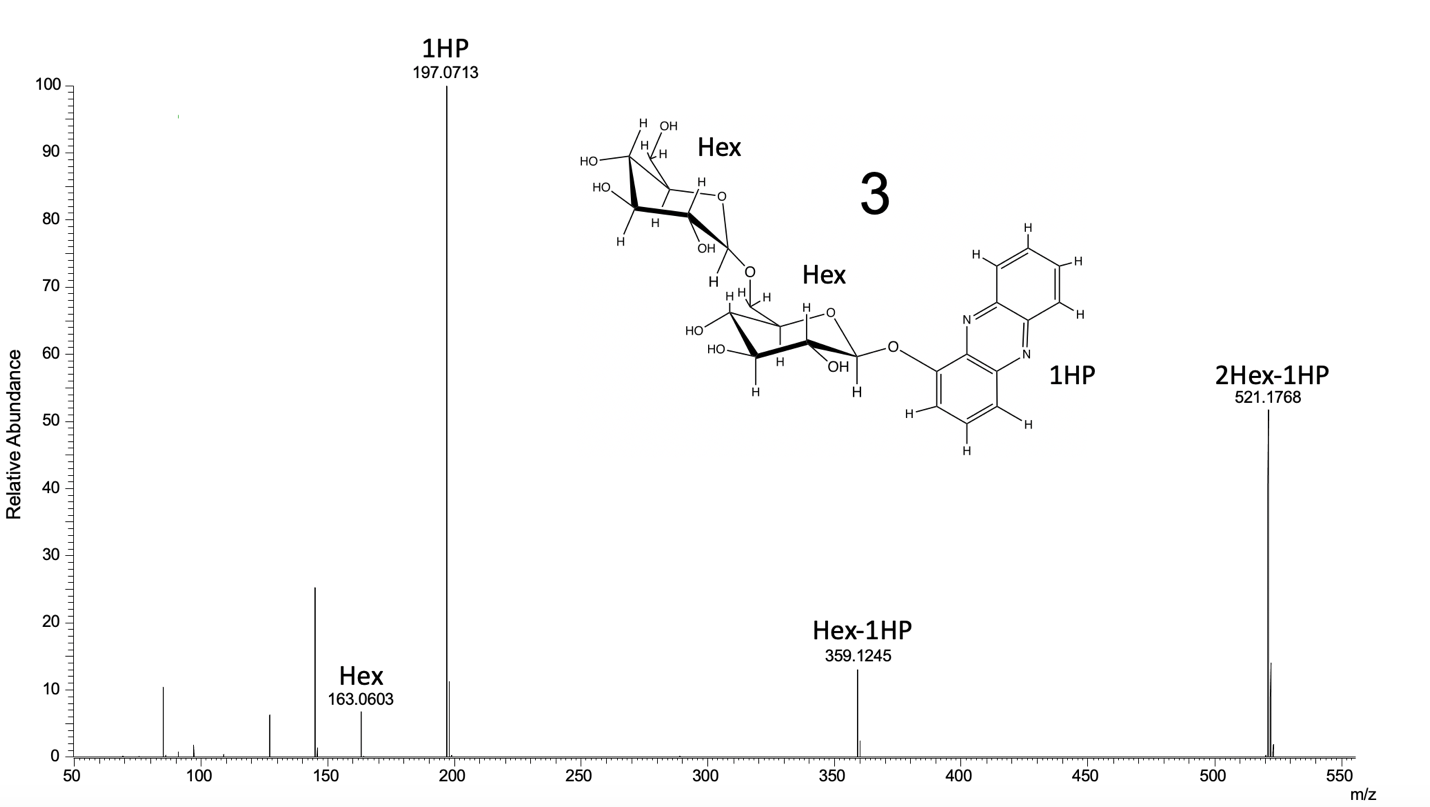

**Supplemental Figure 4.** MS/MS data for (3).

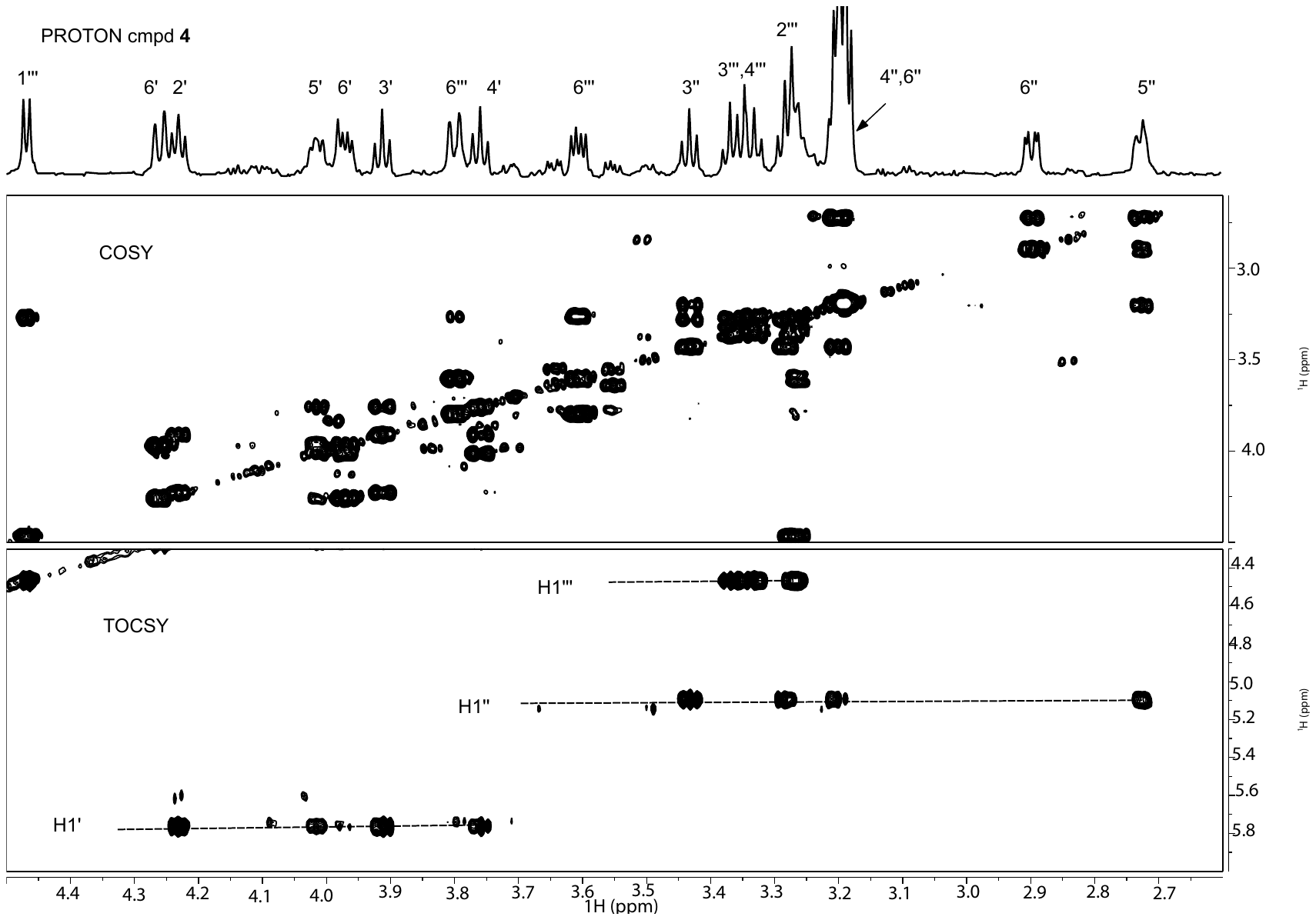

**Supplementary Figure 5.1**: Proton NMR assignments of compound (**4)** glucosyl residues.

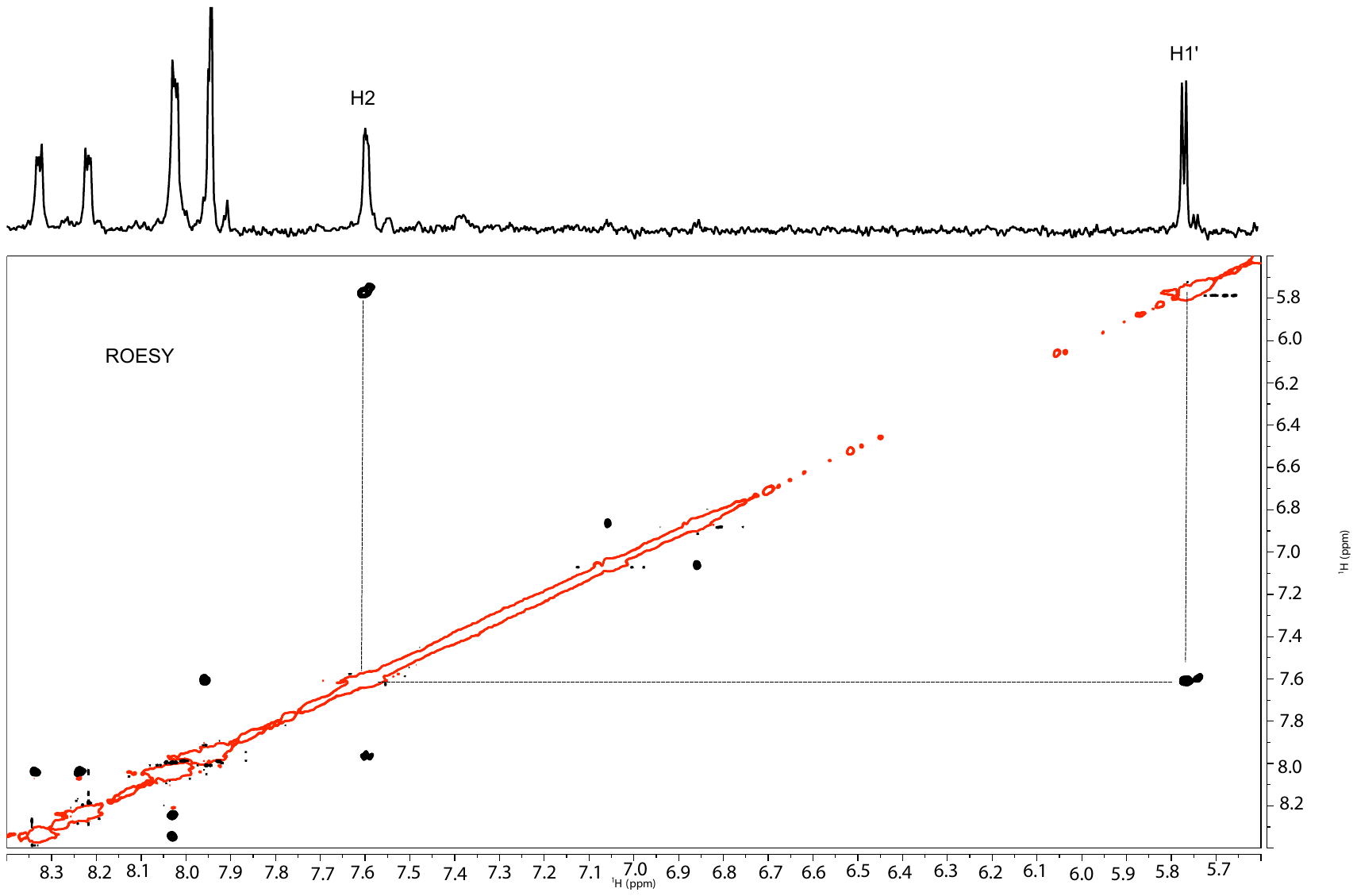

**Supplementary Figure 5.2:** Top panel – the region of 1D proton of compound (**4**) showing phenazine protons and H1’. Bottom panel – the region of 2D ROESY spectrum showing NOE between H2 of phenazine and H1’ of the glucosyl residue linked to position 1 of phenazine.

**Supplemental Table 5.3 (A):** NMR data table showing assignments of Compound (**4**).

**A**

| **#** | **Type** | **d_H_ [ppm]** | **J (Hz)** |
| --- | --- | --- | --- |
| 2 | dd | 7.59 | 6.02, 2.94 |
| 3-4 | dd | 7.94 |  |
| 6 | ddd | 8.33 |  |
| 7-8 | ddd | 8.02 |  |
| 9 | ddd | 8.21 |  |
| 1’ | d | 5.76 | 7.59 |
| 2’ | t | 4.23 | 8.16 |
| 3’ | t | 3.91 | 8.26 |
| 4’ | t | 3.76 | 10.46 |
| 5’ | t | 4.01 | 8.44 |
| 6a’ | m | 3.97 |  |
| 6b’ | d | 4.26 | 11.21 |
| 1’’ | d | 5.1 | 7.68 |
| 3’’ | t | 3.43 | 9.3 |
| 4’’ | m | 3.19 |  |
| 5’’ | t | 2.73 |  |
| 6a’’ | dd | 2.89 | 12.3, 4.22 |
| 6b’’ | m | 3.19 |  |
| 1’’’ | d | 4.47 | 7.71 |
| 2’’’ | m | 3.27 |  |
| 3’’’ | m | 3.35 |  |
| 4’’’ | m | 3.35 |  |
| 6a’’’ | dd | 3.61 | 6.21, 12.41 |
| 6b’’’ | d | 3.8 | 11.95 |

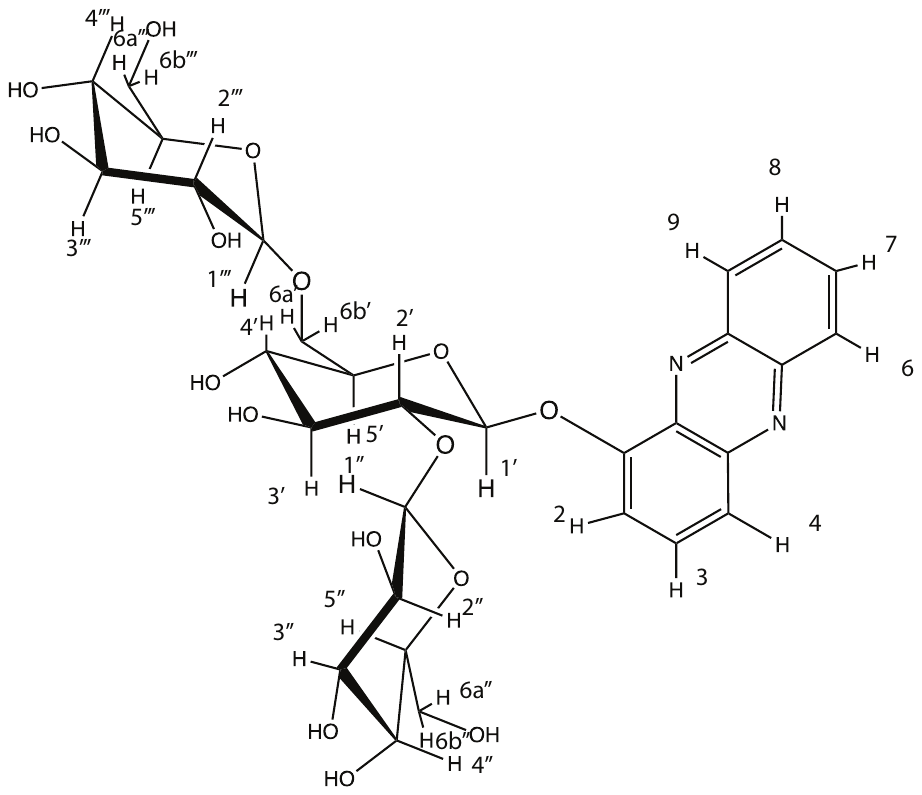

**B**

**Supplemental Figure 5.3:** NMR data of compound (**4**) in D_2_O. (A) Table showing assignments. d indicates a doublet, dd indicates a doublet of doublets, ddd indicates doublet of doublet of doublets, t indicates a triplet, and m indicates a multiplet. (B) Structure of compound (**4**).

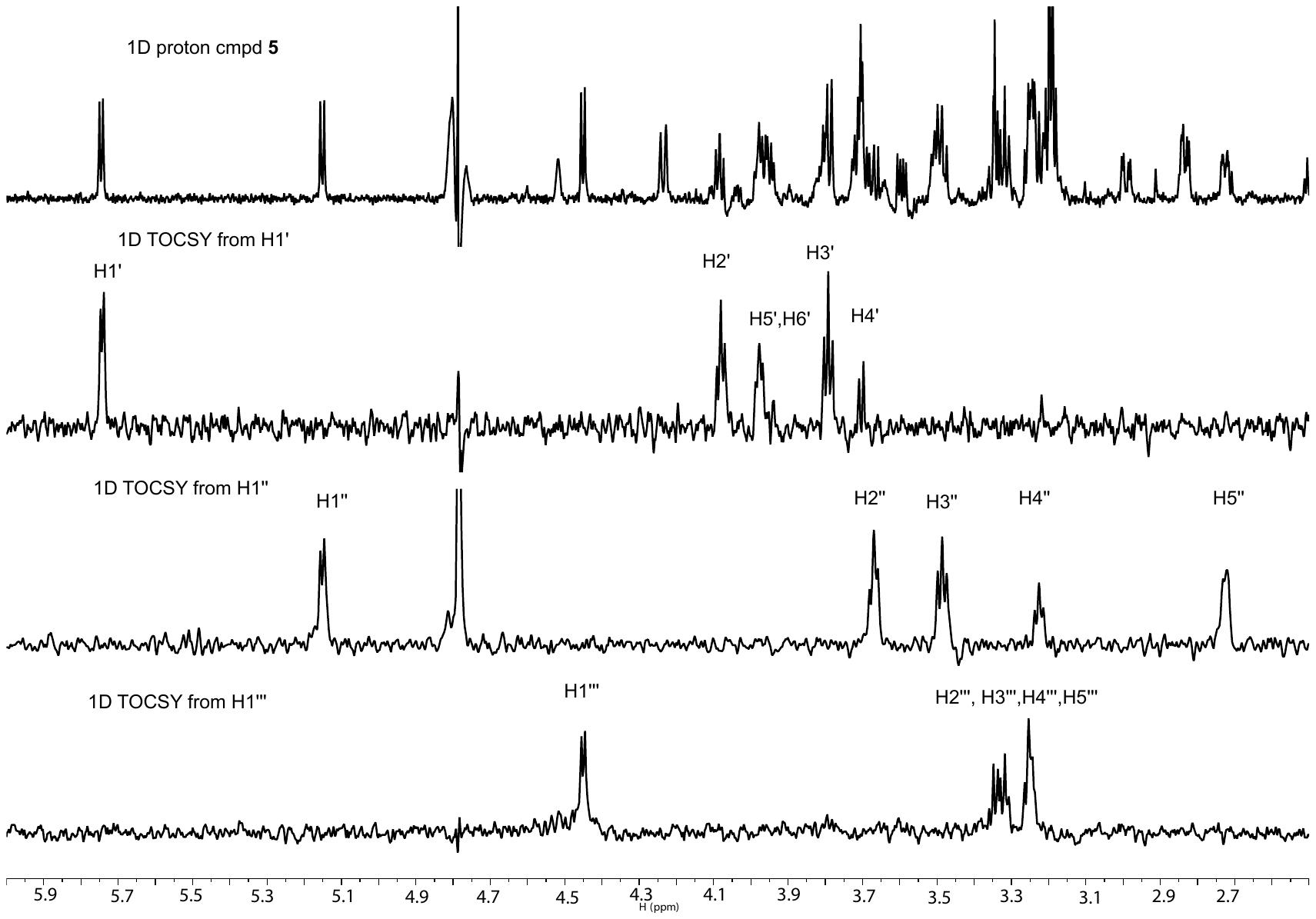

**Supplementary Figure 6.1:** Proton NMR spectra of compound (**5**). The three lower panels of 1D TOCSY spectra show signals from H1-H5 for the two glucosyl residues and the NAcetyl-glucosamine residue.

^1^H (ppm)

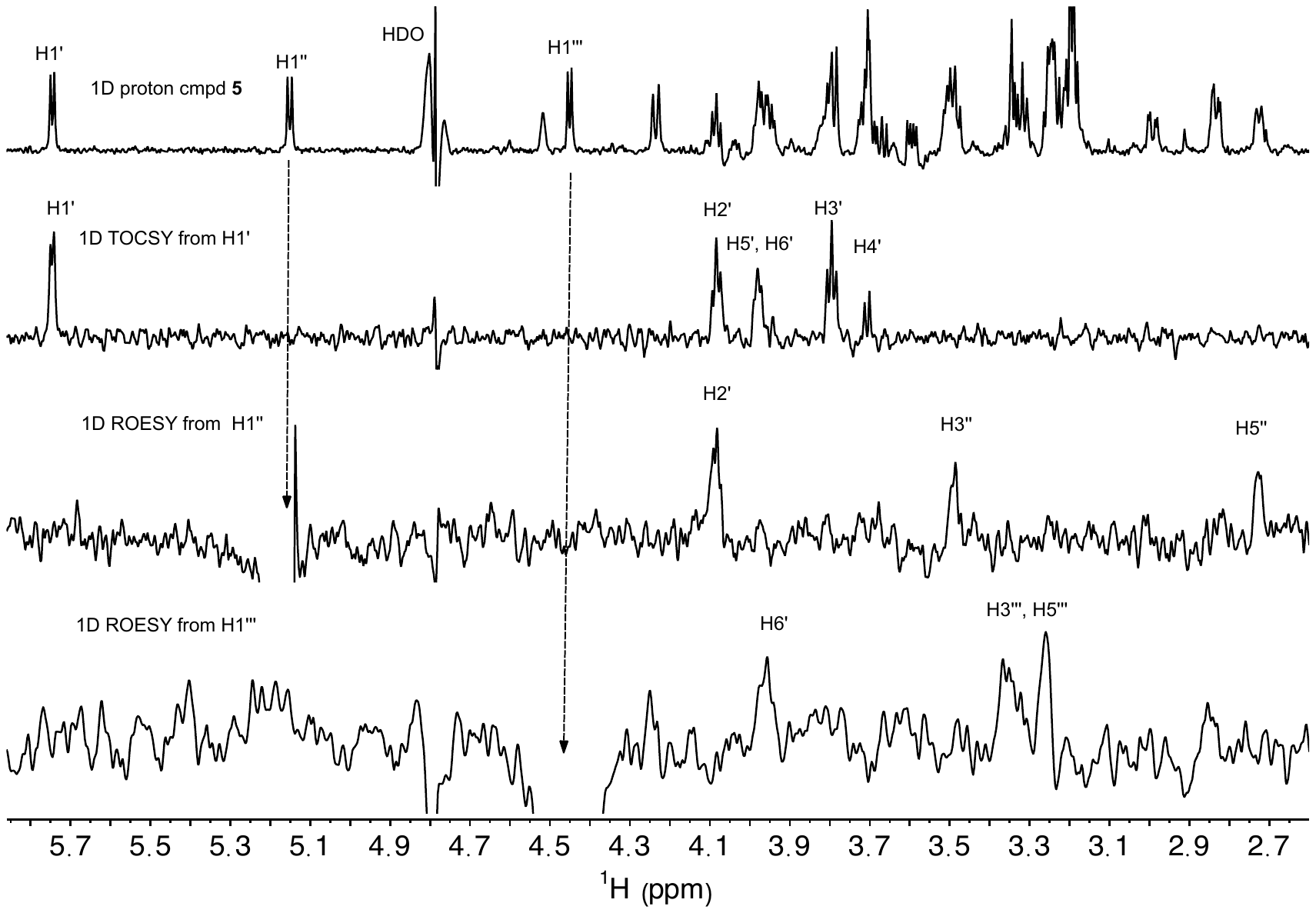

**Supplementary Figure 6.2:** Regions of 1D TOCSY and ROESY spectra of compound (**5**) confirm the terminal glucose and NAcetyl-glucosamine linkages to positions 2’ and 6’ of the glucosyl residue attached to phenazine. The arrows indicate the peaks of interest that have been selected for the 1D ROESY experiment. They correspond with the anomeric protons of each of the sugar residues of the compound.

**Supplemental Table 6.3 (A):** NMR data table showing assignments of Compound (**5**).

**A**

| **#** | **Type** | **δ_H_ [ppm]** | **J (Hz)** |
| --- | --- | --- | --- |
| 2 | dd | 7.59 | 6.29, 1.91 |
| 3-4 | dd | 7.96 |  |
| 6 | ddd | 8.34 |  |
| 7-8 | ddd | 8.04 |  |
| 9 | ddd | 8.26 |  |
| 1’ | d | 5.75 | 7.41 |
| 2’ | t | 4.08 | 8.16 |
| 3’ | m | 3.79 |  |
| 4’ | m | 3.68 |  |
| 5’ | m | 3.95 |  |
| 6’ | m | 3.96 |  |
| 1’’ | d | 5.15 | 8.32 |
| 2’’ | dd | 3.66 |  |
| 3’’ | m | 3.49 |  |
| 4’’ | m | 3.24 |  |
| 5’’ | ddd | 2.72 |  |
| 1’’’ | d | 4.45 | 7.7 |
| 2’’’ | m | 3.33 |  |
| 3’’’ | m | 3.33 |  |
| 4’’’ | m | 3.33 |  |
| 5’’’ | m | 3.24 |  |

**B**

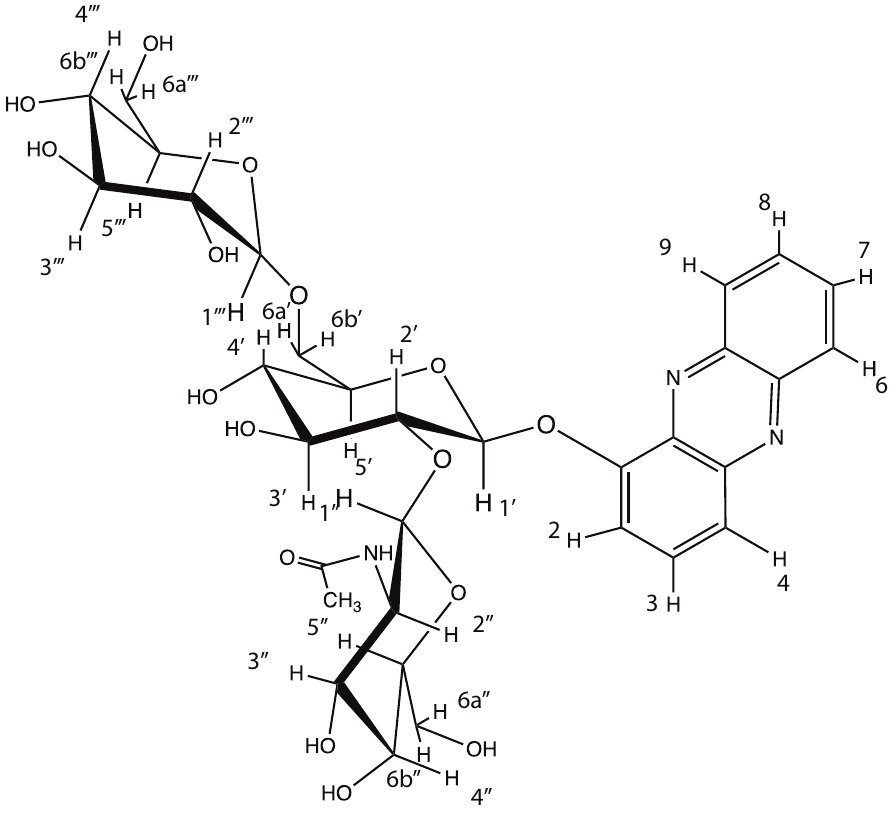

**Supplemental Figure 6.3:** NMR data of compound (**5**) in D_2_O. (A) Table showing assignments. d indicates a doublet, dd indicates a doublet of doublets, ddd indicates doublet of doublet of doublets, t indicates a triplet, and m indicates a multiplet. (B) Structure of compound (**5**).

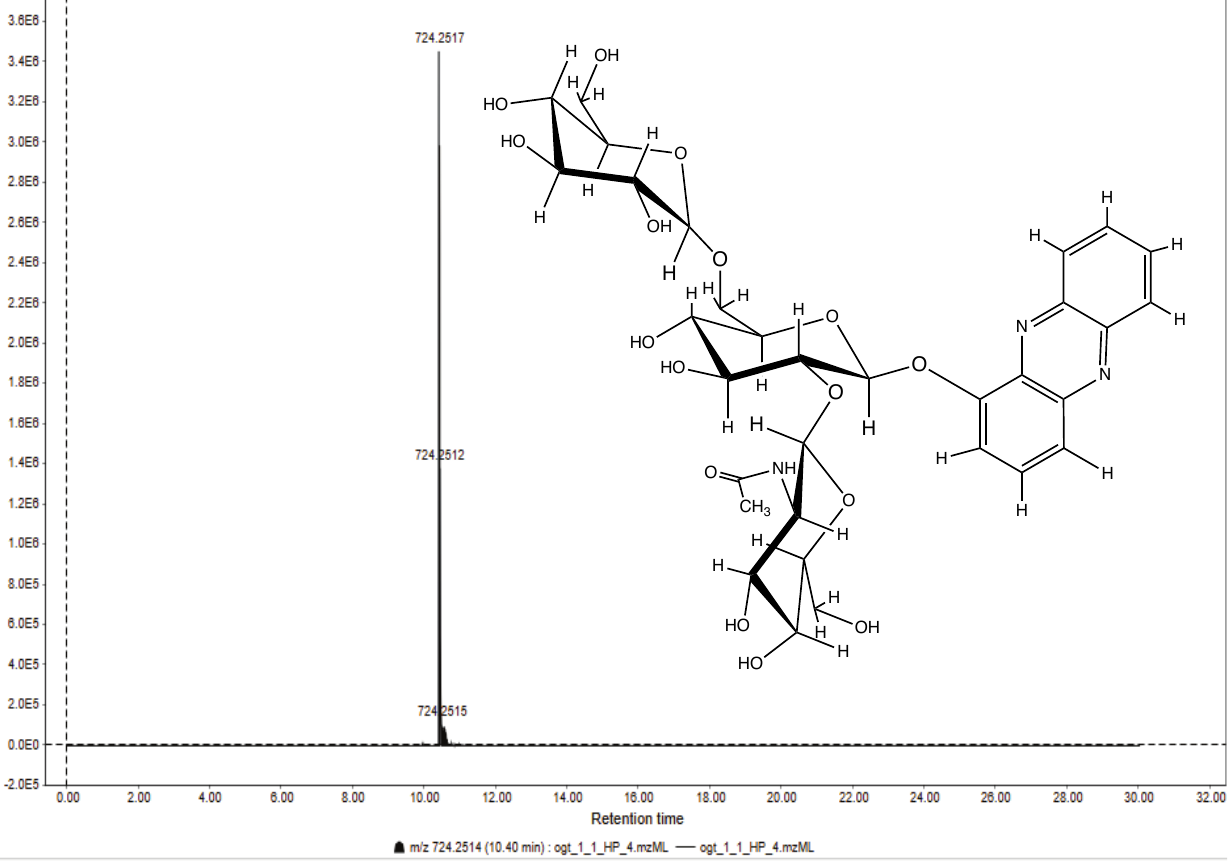

**Supplemental Figure 7.** Selected ion chromatogram for Compound (**5**) in the *ogt-1* knockout mutant exposed to 22.3 μM 1-HP.
